# Task-Related EEG Source Localization via Graph Regularized Low-Rank Representation Model

**DOI:** 10.1101/246579

**Authors:** Feng Liu, Jay Rosenberger, Jing Qin, Yifei Lou, Shouyi Wang

**Affiliations:** Department of Industrial and Manufacturing Systems Engineering University of Texas at Arlington Arlington, TX, 76019, USA; Department of Mathematical Sciences Montana State University, Bozeman, MT, 59717, USA; Department of Mathematical Sciences University of Texas at Dallas, Richardson, TX, 75080, USA

**Keywords:** EEG Source Imaging, Low-Rank Representation, Graph Regularization, Alternating Direction Method of Multiplier (ADMM)

## Abstract

To infer brain source activation patterns under different cognitive tasks is an integral step to understand how our brain works. Traditional electroencephalogram (EEG) Source Imaging (ESI) methods usually do not distinguish task-related and spurious non-task-related sources that jointly generate EEG signals, which inevitably yield misleading reconstructed activation patterns. In this research, we argue that the task-related source signal intrinsically has a low-rank property, which is exploited to to infer the true task-related EEG sources location. Although the true task-related source signal is sparse and low-rank, the contribution of spurious sources scattering over the source space with intermittent activation patterns makes the actual source space lose the low-rank property. To reconstruct a low-rank true source, we propose a novel ESI model that involves a spatial low-rank representation and a temporal Laplacian graph regularization, the latter of which guarantees the temporal smoothness of the source signal and eliminate the spurious ones. To solve the proposed model, an augmented Lagrangian objective function is formulated and an algorithm in the framework of alternating direction method of multipliers is proposed. Numerical results illustrate the effectiveness of the proposed method in terms of reconstruction accuracy with high effciency.

## 1. Introduction

As a direct measurement modality of neural electrical firing patterns, electroencephalogram (EEG) has a higher temporal resolution up to millisecond compared to positron emission topography (PET) and functional magnetic resonance imaging (fMRI) [1–3]. However, a limitation of EEG is its low spatial resolution since the measurement is on the scalp rather than inside the brain. Given the recorded scalp EEG data, to reconstruct of the brain sources signal inside the brain is known as EEG inverse problem or EEG source imaging (ESI) [4]. ESI technique has been widely used in the study of language mechanisms, cognition process, sensory function, as well as the localization of epileptic seizure [1, 5]. ESI is also an preliminary step for brain connectivity analysis [6, 7] on the source level [8–11].

Since the number of electrodes usually outnumbers that of brain sources, the ESI problem is highly ill-posed or mathematically underdetermined. A variety of methods have been proposed to address this challenging problem with different neurophysiological assumptions,formulated by various regularization techniques [4, 12]. There are two main types of inverse solvers for ESI: dipole fitting and distributed inverse solvers [13]. Dipole fitting empirically solves the MEG/EEG forward and inverse problems by characterizing the neural generators responsible for electrical potential detected on the scalp sensors [4, 14]. The algorithms proposed in recent years to solve the ESI problem within distributed dipoles paradigm can be further summarized into three categories in general, which are the (1) Bayesian framework [15–20], (2) state-space based algorithms [21–25], (3) models using sparse representation technique [13, 26 -31].

In many sensory or cognitive studies, the underlying activated cortical responsible for the signal processing are relatively focal and thus sparse, which makes the spatial sparse constrain not only mathematically necessary but also neurologically reasonable [1]. One widely accepted assumption is that a sparse spatial structure is favored than a complicated source configuration to explain the same data [32]. An representative early pioneering work is the *ℓ*_2_-norm based minimum norm estimate (MNE) inverse solver [33]. Based on MNE algorith, Pascual-Marqui et al. later proposed standardized low-resolution brain electromagnetic tomography (sLORETA) [34] that enforces spatial smoothness of the neighboring sources and normalizes the solution with respect to the estimated noise level. Some algorithms proposed to use combined multiple solvers, e.g., Weighted minimum norm-LORETA (WMN-LORETA) which combines the LORETA solver and a weighted minimum norm to compensate for deeper sources originate from the subcortical regions[35]. As the above mentioned algorithms are based on *ℓ*_2_-norm to different extents, the estimated source area is over-diffuse. By replacing *ℓ*_2_-norm by *ℓ*_1_-norm, minimum current estimate (MCE) [36] is proposed to overcome overestimation of active area sizes incurred by *ℓ*_2_-norm. Recent development of compressive sensing algorithm proved the *ℓ_p_* (*p* ≤ 1) regularization on the original source signal usually provides a set of discrete sources distributed across the cortex due to high coherence of lead field matrix, in order to encourage reconstruction of extended source patches, it has been found that by enforcing sparsity in a transformed domain, e.g., total variation (TV) regularization [29, 37–39], focal source extents can be better estimated.

The aforementioned algorithms estimate source location at each time point independently, leading to discrepancy along the time direction. To encourage temporal smoothness, a number of regularization techniques based on spatiotemporal mixed norms have been developed, including the famous Mixed Norm Estimates (MxNE) which uses *ℓ*_1,2_-norm regularization [27], and time-frequency mixed-norm estimate (TF-MxNE) which uses structured sparse priors in time-frequency domain for better estimation of the non-stationary and transient source signal [26]. The disadvantage of MxNE is imposing *ℓ*_2_ norm can not guarantee the temporal smoothness in neighboring time points when the source signal is corrupted, while the limitation of TF-MxNE is the lost of high temporal resolution by using of STFT Gabor matrix as a dictionary for time-frequency coefficients. Due to the existence strong spontaneous background source activations, discriminative source activation pattern corresponding to different cognitive tasks which provide more insights shall be reconstructed, Liu et al proposed to use label information of EEG data to find those discriminative source activation pattern [40, 41].

One common drawback of the existing ESI algorithms is that they only consider noises on the sensor level and ignore the spurious noise from the cortex. If perfectly reconstructed, the estimated source is aggregated by task-related source and spurious noise in the source space. The true task-related sources will be corrupted by spurious sources, which motivates us to develop new algorithms to find true task-related source. There are two commonly accepted assumptions (1) spatially sparse (2) temporally continuous for the task-related source activation pattern, which inevitably leads to the low-rank property of the source space. To better discover the task-related source, we impose the low-rank term in the goal function as we consider it is a more direct constraint for spatial sparse. Also, we use a more direct penalty term to temporal smoothness, which is to penalize dissimilarity of temporally neighboring samples based on manifold graph embedding. It is worth-noting that we used the graph regularization term in our previous paper, however the graph is defined to be fully connected for all the points within one class [40], which inevitably drive all the activate patterns at different time points having the same magnitude, thus making the previously defined graph regularization term rely on a strong assumption and limit its future application for realistic cases.

In this paper, we propose a novel EEG source imaging model based on temporal graph regularized low-rank representation. The model is solved based on the alternating direction method of multipliers (ADMM) [42]. We conducted extensive numerical experiments to verify the effectiveness on discovering task related low-rank sources. The reconstructed solution is temporally smooth and spatially sparse. The contributions of our paper are summarized as follows:

1. We propose to consider the noise not only in the sensor level, but also in the source space.
2. A low-rank representation model (LRR) is proposed for the first time on EEG inverse problem inspired by the low-rank property of true task-related source configurations.
3. We redefined graph embedding regularization based on our previous paper that utilizes temporal vicinity information of samples to promote temporal smoothness.
4. A new algorithm based on ADMM is given which is capable of extracting the low-rank task-related source patterns.

## 2. Inverse Problem and Temporal Graph Structures

In this section, we briefly review the inverse problem and then discuss the design of temporal graph regularization.

### 2.1. The Inverse Problem

The cortex source activations propagate to EEG sensors through a linear mapping matrix called lead field matrix, and it can be described as the following linear model:

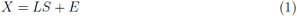

where 
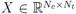
 is the EEG data measured at a set of *N_c_* electrodes for *N_t_* time points, 
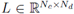
 is the lead field matrix which maps the source signal to sensors on the scalp, each column of *ℓ* represents the electrical field of a particular source to the EEG electrodes, 
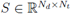
 represents the corresponding driving potential in *N_d_* sources locations for the *N_t_* time instants. Since the number of sources is much larger than electrodes, solving *S* given *X* is ill-posed with infinite feasible solutions, which necessitates a regularization term to be imposed. Generally, an estimate of *S* can be found by minimizing the following cost function, which is composed of a quadratic error and a regularization term:

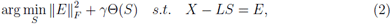

where ‖.‖_*F*_ is the Frobenius Norm. The penalty term Θ(*S*) is to encourage neuro-physiologically plausible solutions. The regularization term take the form of *ℓ*_2_, *ℓ*_1_ or mixed norm. For example, spatially smooth formulation as in LORETA estimation or spatially sparse formulation with Least Absolute Shrinkage and Selection Operator (LASSO) estimate.

### 2.2. Temporal Graph Embedding

An important assumption on the source signal is that two temporal adjoint data points should have similar intrinsic activation pattern. In computer vision community, a lot of manifold learning methods have been proposed to find intrinsic similar structure on low-dimensional sub-manifolds embedded in a high dimensional ambient space, such as locally linear embedding [43], Locality Preserving Projection [44], Neighborhood Preserving Embedding [45]. A graph can be viewed as geometric neighborhood relationship between each vertex representing each data sample, the weight between vertex represents similarity between two points [46]. Inspired by the manifold theory [47] and work from Liu et al [48], we use a regularization term to penalize the difference of two neighboring source signal. In our previous work, we use a graph regularization term to promote intra-class consistency [40], but the assumption is too strong by requiring all the reconstructed sources at different time points has the same location as well as signal magnitude as long as they belong to the same class. Now define a temporal graph regularization as

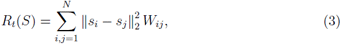

where *s_i_* is the *i*-th column of the matrix *S*, and a binary matrix *W* is designed as follows

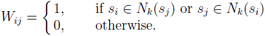

The graph embedding matrix *W* contains temporal vicinity information. *N_k_*(*s_i_*) is the set containing *k* temporally closest points to *s_i_*. This formulation intends to force neighboring source signal having similar pattern. The benefits are twofold, one is for temporal smoothness, another advantage is to make the reconstructed source denoised for intermittent spurious source activates. By defining *D* as a diagonal matrix whose entries are row sums of the symmetric matrix *W*, i.e., *D_ii_* =∑_*j*_*W_ij_*, and denoting *G = D-W*, *R_t_(S)* can be rewritten as:

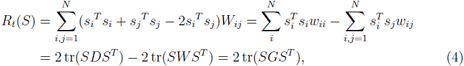

where tr(·) is the trace operator of a matrix, i.e., adding up all diagonal entries of a matrix.

## 3. Proposed EEG Source Imaging Model

Before we present our low-rank model with temporal graph structures, we comment on the limitations of traditional model and come up with the decomposition of task-related source with low-rank property and spontaneous non-task-related spurious sources that is sparsely distributed spatially and with transient patterns. We come up with a graph regularized low-rank representation model and detailed discussion on the purpose of each term in the goal function.

**Figure 1:**
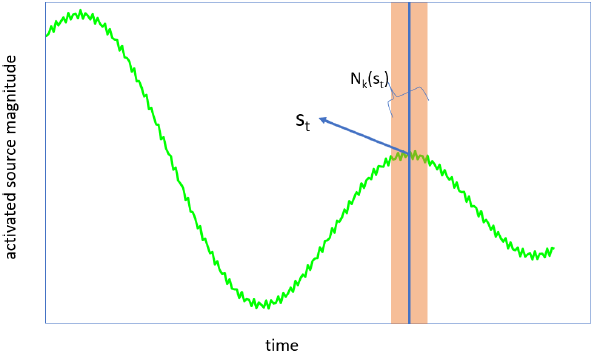
Illustration of temporal smoothness. By design the temporal graph matrix G, the reconstructed signal should have consistent pattern within the same neighborhood window.

### 3.1. Decomposition of True and Spurious Sources

In general, two types of noises should be considered, one originates from inaccurate measurement of the sensors modeled by Guassian white noise, which is denoted as *E* in Eq.(1), the other type of noise is called biological noise that comes directly from the spontaneous activations in the source space, which are not task-related and termed as spurious source. The second types of noise (spurious source) also contributes to the EEG signal in the same way as ground truth source. A drawback of traditional models is that they didn’t distinguish the spurious sources from the true sources, and the estimated source can be composed of both task-rated source and spurious sources. To address the above mentioned problem, we propose to use a decomposed source spaces, composed of a low-rank source space and spurious sources originates from spontaneous biological noises. The illustration of decomposition of source space as well as the whole procedure is given in Fig.2, where *S*_1_ has a low-rank property and *S*_2_ is sparse, and sum of *S*_1_ and *S*_2_ is no long low-rank, making *X* lose low-rank structure.

**Figure 2:**
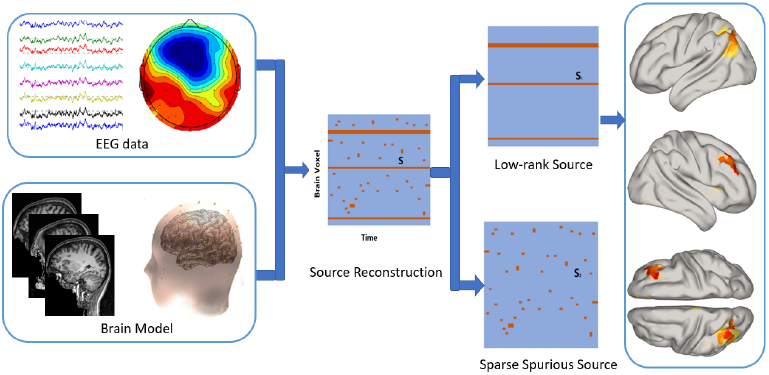
Extraction of low-rank true source from spurious source pipeline: After gathering the MRI scans of the head, tissue segmentation is conducted followed by mesh generation. By assigning conductivity values to different tissues and electrodes co-registered with the meshing model, boundary element method (BEM) was used to solve the forward model. Each triangle represents a brain source, the direction of the source is assumed to be perpendicular to the triangular surface. With EEG data and forward brain model, source reconstruction is calculated. The factual source signal *S* can be decomposed into two source matrix. The task related true sources S_1_ have a low-rank property and the spurious sources S_2_ are the sparse but not temporally consistent. The low-rank source solution is projected to cortex voxels to illustrate the activation pattern.

### 3.2. Basic Low-Rank Representation (LRR) Model

We argue that during a short period of task-related Evoked Repose Potential (EPR), the number of corresponding activated sources is sparse and remain activated during this period of time, which makes the EPR source matrix low-rank with most rows being zero. However, the brain signal is known to be heavy noisy, spontaneous activations are also active from different source spots. The low-rank representation model for the EEG inverse problem is introduced as follows:

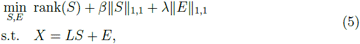

where *β*, λ and γ are positive scalars to balance the rank function, sparsity cost of sources and the reconstruction error. It is pointed out that *ℓ*_1_ on the error term is more robust to outliers, we use *ℓ*_1_ in the row-rank model [49]. The ‖*E*‖_1,1_ is defined as

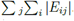
. Due to the discrete nature of the rank function, it is a common practice to use a surrogate nuclear norm ‖.‖_*_ instead. The goal function is given as:

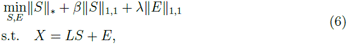

The above formulation is a simple version trying to estimate the task-related source activation pattern by using low-rank constraint. To promote the temporal smoothness, the Laplacian graph structure is included in the next section followed by the optimization algorithm.

### 3.3. LRR Model with Graph Regularization

Incorporating the previous temporal graph structures, we introduce our proposed model called low-rank representation with temporal graph structures ESI (LRR-TG-ESI). The model is composed of data fitting term to explain the EEG data, temporal graph embedding regularization term that promote temporal smooth, and *ℓ*_1_ norm for sparsity penalty and nuclear norm for the low-rank structure of ground true source. By combining the low-rank prior and the temporal graph regularization, we propose the following model for ESI:

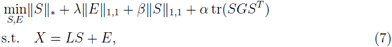

where λ,*β*,α > 0 are tuning parameters to balance the trade-off of different terms. Our proposed model is able to enforce row-sparse via low-rank prior and temporal smoothness via temporal graph regularization while fitting the EEG data *X*. Compared to earlier works on the ESI problem, both the low-rank prior and graph regularization is novel, although the graph regularization term has been discussed in our early paper [40], but it is not defined on the temporal manifold, and the previous definition in [40] make the magnitude of source signal to be equal intra-class, which is not realistic in real world. To further consider the spatial smoothness, a total variation term can be imposed as another penalty term, such as first order total variation (TV) regularization in Ref.[38, 50], fractional order TV in [37, 39], and similar algorithm can be derived under the framework of ADMM, however further investigation with constraints of TV is our future work.

## 4. Numerical Algorithm

To solve (7), an algorithm in the ADMM framework is developed. The augmented Lagrangian function of (7) is

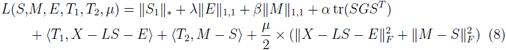

By some simple algebra, (8) can be reformulated as

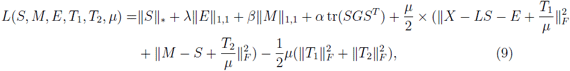

where *T*_1_ and *T*_2_ are Lagrangian multipliers and *μ* is a parameter for the augmented Lagragian term. The variables are updates alternately in a Gauss-Seidel manner by minimizing the augmented Lagrangian function, with other variables fixed. For symbolic simplicity, we rewrite Eq.(9) into the following form:

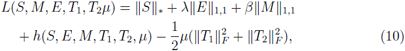

where

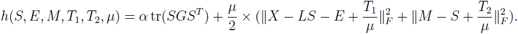

If the augmented Lagrangian function is difficult to minimize with respect to a variable, a linearized approximate surrogate function can used, hence the algorithm bears the name Linearized Alternating Direction method [46, 51]. Updating *S* by minimizing 
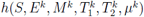
 (suppose we are at iteration *k*) is equivalent to minimize the following goal function with the other variables fixed:

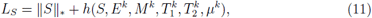

which is approximated by optimizing its linearizion at 
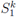
 plus a quadratic proximal term, given as:

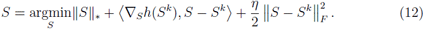

Here *η* is a constant satisfying

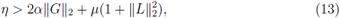

where ‖.‖_2_ is the spectral norm of a matrix, i.e, the largest singular value. As long as (13) is satisfied, (12) is a good approximate to (11). The solution to (12 has a closed form using a singular value thresholding operator (SVT) [52] given as:

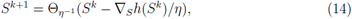

where Θ_ε_(A) = *USε(∑)V^T
^* is the SVT operator, in which *U∑V^T^* value decomposition of A and *S_ε_(s)* is defined as 
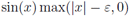

is the singular 
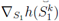
is calculated as

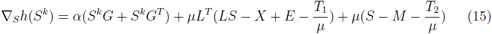

To update *M* and *E*, it is equivalent to solve the following problem:

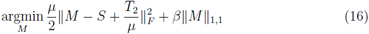

**Algorithm 1.**
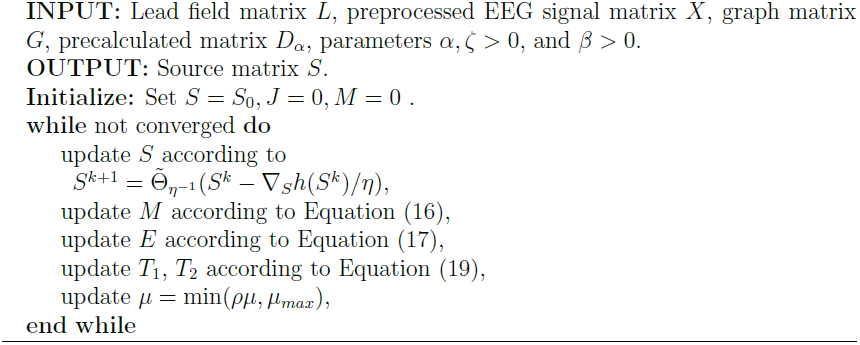
Source Imaging Based on Spatial and Temporal Graph Structures

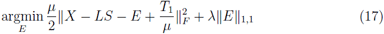

The general form of (16)–(17) is a *ℓ*_1_ norm proximal operator defined as

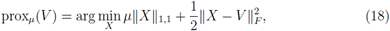

with *μ* > 0. The above problem (18) has a closed form solution, called soft thresholding, defined by a shrinkage function,

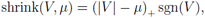

where (*x*)_+_ is *x* when *x* > 0, otherwise 0. The shrinkage function is defined as element-wise operator. Problem (16)–(17) has a close form solution described with the shrinkage function. After updating all the variables, these Lagrange multipliers are updated by

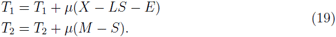

The parameter *μ* is updated by *μ* = min(*ρμ*, *μ*_*max*_). A summarized algorithm is given as Algorithm 1. We initialize the *S* with the estimate *S*_0_ from *ℓ*_1_ solver.

It’s worth noting that the data fitting term we use is *ℓ*_1,1_ norm of *E* in the model, and there are other options. Generally, if the Gaussian noise *E* is small, then the norm ‖*E*‖_F_, is an appropriate choice, but for random data corruption, *ℓ*_1,1_ should be used, and for sample specific data corruption, *ℓ*_2,1_ [53–55], should be used. It has been shown that *ℓ*_2,1_ is more robust to large outliers in some samples. Although the norm used in Algorithm 1 is *ℓ*_1,1_, it can be extended to *ℓ*_2,1_ norm of *E*, where ‖*E*‖_2,1_ is defined as

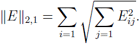

In stead of solving (17) to update *E*, the following goal function (20) needs to be solved to update *E*.

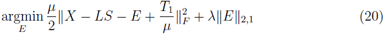

By substituting 
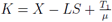
, if *E*^*^ is the optimal solution of

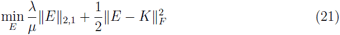

Based on the Lemmas in Ref[56], the solution to (21) is

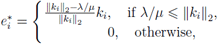

where 
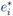
 and *k*_*i*_ is the *i*-th column of matrix *E^*^* and *K* respectively.

**Convergence:** The convergence of Algorithm 1 can be easily derived from [51]. Even though the update of *M* and *E* is separated in Algorithm 1, they can be combined in one step to become a larger block step and simultaneously solving for (*M*, *E*) which is the same case described by LADMAP algorithm [51]. The convergence analysis in [51] can be applied to our case, thus the algorithm convergence is guaranteed [46].

## 5. Numerical Experiments

In this section, we conducted several experiments to illustrate the effectiveness of our proposed method. Since both the nuclear norm and the graph regularization is relative new for the ESI problem, we started from the simple model (6) to help the readers understand the property and impacts of low-rank prior along with data fidelity term and sparsity term for source reconstruction. In the first experiment, we did a comprehensive exploration for different parameter settings by varying the weights between low-rank term, data fitting term and sparsity term. In the second experiment, we illustrate the temporal smoothing functionality of the graph regularization term for uncorrupted smooth source and corrupted source with abrupt signal jumps. In the third experiment, we give comprehensive numerical results by testing our algorithm against the benchmark algorithms to showcase the effectiveness of the proposed method in reconstructing task-related source, where we show that our algorithm can not only find the activated locations, but also reconstruct the time-course of source activation with high precision.

### 5.1. Head Model

A realistic head model, referred to as New York Head model [57], is used in our numerical experiment. The New York Head model is based on highly detailed MRI images derived ICBM152 anatomy, which is a nonlinear average of the T1-weighted structural MRI of 152 adults and calculated with state-of-the-art finite element electrical modeling. The New York Head model is considered to be highly accurate since it considers six tissue types when conducting segmentation, which is scalp, skull, CSF, gray matter, white matter, air cavities, with a native MRI resolution of 0.5 mm^3^. The dimension of the lead field matrix used in the numerical experiment part is 2004 by 108, representing a linear mapping from 2004 sources to 108 electrodes.

### 5.2. Experiment 1: Test on Simple Low-rank Model

To understand the property of low-rankness, we start from a simple model to help the readers understand the property of low-rank in the EEG source imaging and the validity of low-rank when recovering the accurate location as well as time course of source signal. Like in [58], eight octants are divided as regions of interest (ROI) are considered, which are Right Anterior Inferior (RAI), Right Anterior Superior (RAS), Right Posterior Inferior (RPI), Right Posterior Superior (RPS), Left Anterior Inferior (LAI), Left Anterior Superior (LAS), Left Posterior Inferior (LPI), Left Posterior Superior (LPI). In the simple experiments, we selected 2 different ROIs and randomly select one activated voxel in each of these ROIs, and a 4*th* order moving average time series is generated, as illustrated in Fig.3.

**Figure 3:**
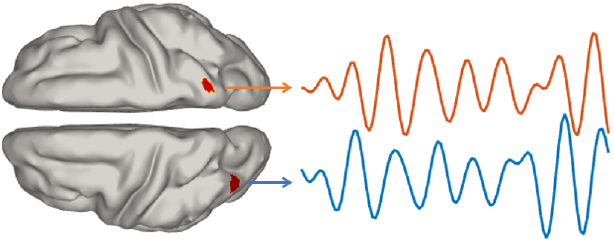
Illustration of two activated sources time series on two different ROIs.

At each location, a time series with length of 500 were generated to represent the source activation time-course. At each time point, two randomly picked sources are activated to simulate the non-task related spurious noise with mean of 0 and variance to be 1. The task-related activate pattern has low-rank property, however, the noise corrupted source space is no longer low-rank. We repeated our experiment 50 times for all the combinations of λ and *β*, where λ = {0.01, 0.02, 0.03, 0.05, 0.1, 0.2, 0.5} and *β* = {0.005, 0.01, 0.015, 0.02, 0.05, 0.1}. The reconstructed error (RE) metric used here is

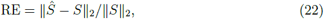

where 
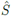
 represents the reconstructed source. We define the 
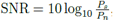
 where *P*_*s*_ and *P*_*n*_ are the power of signal and noise respectively. The violin plot is used to visualize the distribution of reconstruction errors, in corresponding to *β* and λ respectively in Fig.4 and Fig.5. Increasing *β* will penalize the strength of signal and make the reconstruction error to be large when λ is small. However, When λ is set to be 0.5, the weight of data fidelity is high, thus driving the solution have a better data fidelity, which can balance off the increase of *β*.

**Figure 4:**
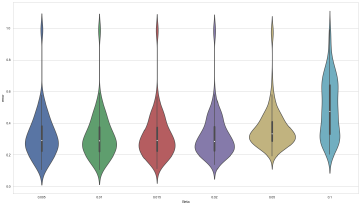
Reconstruction errors varying *β*.

**Figure 5:**
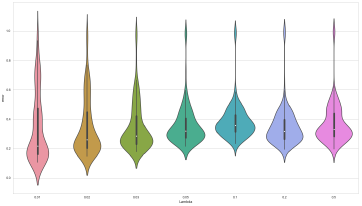
Reconstruction errors varying λ.

The averaged reconstruction error of 50 experiments for all of *β* and λ is given in Fig.6a and the averaged rank for the estimated source for all the combination of *β* and λ is given in Fig.6b. As we increase the value of λ, the rank is also increasing, which underlies the trade-off between explaining the data and finding the latent low-rank structure of the source space. Increasing λ means more weight on the data fitting term, and the spurious source can also be recovered, thus increasing the rank of the reconstructed source.

**Figure 6:**
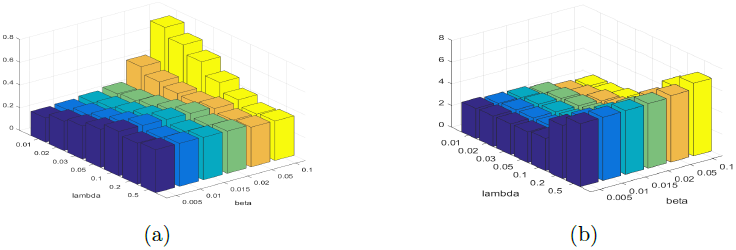
Averaged reconstruction error and rank varying λ and *β* over 50 experiments.(a) average of reconstruction error for different λ and *β* (b) average of rank for the source matrix *S*. Increasing λ means more weight on data fidelity and the rank becomes higher. Our model works well with a wide range of parameter setting.

To empirically understand and explain the reconstruction error in Fig.6a, we visualize the curve fitting performance for truth source time series and the reconstructed source time-course corresponding to different level of reconstruction error. We picked the curve fitting cases when the reconstructed error is 0.2, 0.4, 0.8 and 1 respectively. For error equal to 0.2, we picked one experiment and plot the fitting of time series which demonstrated a very good curve fitting of the ground truth vs the reconstructed one as is shown in part (a) of Fig.7. Also we picked another experiment whose error equal to 0.4 and it is shown in part (b) of Fig.7. The curve fitting is slightly worse than the previous one but it is hard to notice the difference compared to the previous one, however according to our error metric, the error is 0.4. One thing to notice is that in both situations, the reconstructed rank equal to the ground truth rank, which is equal to 2, moreover, the estimated two active source location is exactly the ground true locations. We also examined the case when error is up to 0.8, and the curve fitting plot is given in part (c) of Fig.7. We examined this happened when we set λ = 0.01 and *β* = 0.1. In this case, the penalty for sparsity is 0.1, which means too much penalty for the sparse term and the signal magnitude is reduced by the shrinkage operation. We also notice than in the experiment, the error can be up to 1 no matter what parameter settings are given, as can be seen in the top region of Fig.4 and Fig.5. The curve fitting plot for the failed case is given in part (d) of Fig.7.

**Figure 7:**
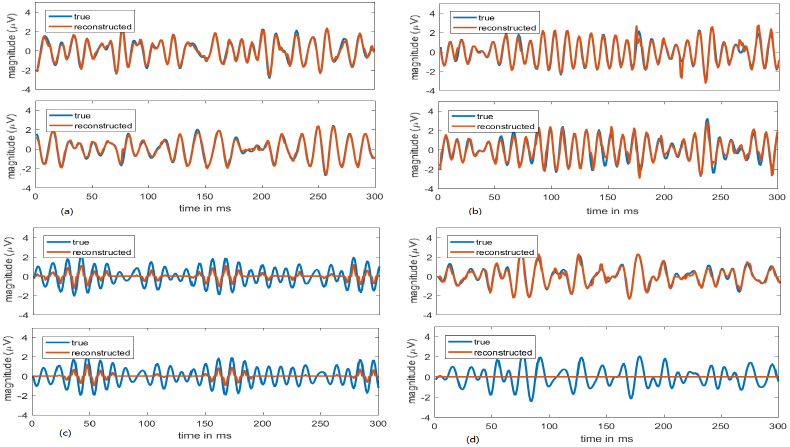
Source time course fitting illustration: (a): ground truth time course vs reconstructed for two activated source at different ROIs in one experiment when fitting error equal to 0.2. Here λ = 0.02 and *β* = 0.005.(b): ground truth time course vs reconstructed for two activated source when fitting error equal to 0.4. Here λ = 0.5, *β* = 0.01. (c): ground truth time course vs reconstructed for two activated source when fitting error equal to 0.8. Here λ = 0.01 and *β* = 0.1. (d): ground truth time course vs reconstructed for two activated source when fitting error equal to 1. Here λ = 0.02 and *β* = 0.01. The curve fitting of (b) is slightly worse than (a), corresponding to the RE= 0.2 and RE= 0.4. When the sparsity parameter is set too large, the reconstructed magnitude is smaller than the ground truth as is shown in (c). For some cases, only the time course in one source location is reconstructed shown in (d) with RE to be 1.0.

Although the situation is very rare when *RE* = 1, we want to visualize the activation pattern on the cortex to see how discrepant the reconstructed location compared to the true source location. For comparison, Fig. 8 is given when *RE* = 0.4 and *RE* = 1. The reconstructed source locations on two ROIs are exactly the same with the ground true location when *RE* = 0.4, when *RE* = 1, one source location is reconstructed perfectly while the other source location is not accurately located, however the neighboring sources are reconstructed.

**Figure 8:**
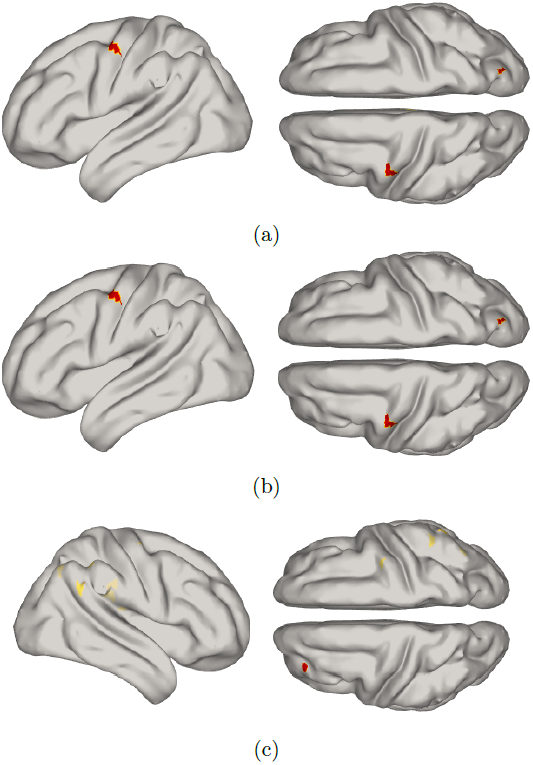
(a) Ground truth source activation pattern (b) Reconstructed source activation when 0.4. This plot illustrates the perfect localization of ground true sources. (c) Reconstructed source activation when *RE* is 1. This plot illustrates when our algorithm failed to recover one of the the exact locations of two activated sources, while the other one can be recovered perfectly. the pictures on the left is the reconstructed location, but still close to the ground truth.

To check how the rank of *S* evolve during the iterations, the boxplot of the rank at selected steps are given in Fig.9. Starting from an initial value with high rank, the rank of 
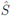
 is decreasing as the iteration proceeds. We set the maximum rank to be 20, during the iteration process, the rank of *S* is converged to very small number for most of the cases.

**Figure 9:**
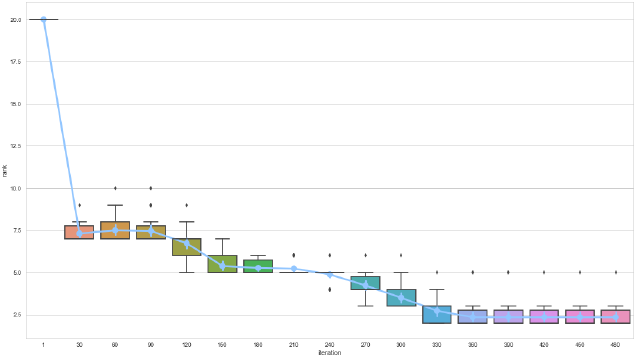
Convergence of rank of *S* during iteration procedure: In the first iteration, we set the maximum rank to be 20, and in most case the rank will converge to 2.

### 5.3. Experiments 2: Test LRR with Temoral Graph Prior

In this section, we solve the LRR-TG-ESI problem (7) with graph regularization term to test its impact on the reconstructed signal. Under the same setting with Experiment 1, we assign different values {0.01, 0.02, 0.05, 0.1, 0.5} for the graph regularization parameter α. The original source signal was smooth, then it was corrupted with randomly number at some time points. There are also 2 randomly picked activated sources representing spurious sources with mean of 0 and variance to be 1. The “temporal smoothing” impact of the graph regularization is shown in Fig.10, where λ = 0.02 and *β* = 0.01. In Fig.10, the original signal is corrupted and not smooth at some time points, we set the neighbor size to be 2 (the closest signal before and after the one to be estimated) when calculating the Laplacian matrix. It is evident from the formulation (3) that the graph regularization term will decrease the dissimilarity of temporally neighbored reconstructed source. If α is set to be 0:5, the graph regularization term penalized heavily on the curvature of the reconstructed signal as is illustrated in Fig.10. We can see that with the temporal graph prior, the reconstructed source is more smooth. It is worth noticing that the main purpose of temporal graph prior is not to smooth the time course for the activated locations, the main purpose is to filter out the spurious activations that are short transients with abrupt jumps. Combined with the low-rank prior, the temporal graph prior can filter the spurious activations and reconstruct the task related activated source. The randomly planted spurious sources are filter out by penalizing the graph regularization and nuclear norm, and in most of the cases, the final rank is 2 can be achieved within a wide range of parameters. The time series plots of original EEG signal, corrupted EEG signal, and EEG signal recovered from the reconstructed sources from (7) by setting λ = 0.02 and *β* = 0.01, as well as the topoplots at 42 ms is given in Fig.11. To illustrate again the impact of the graph regularization, *α* is chosen from {0.01, 0.02, 0.05, 0.1, 0.5}. The first row is the EEG data generated from the task-related sources (persistent and low-rank), the second row is the EEG data from the task-related sources and the spurious source, the SNR is −0.72 dB, which means the energy of spurious source is slightly larger than the task-related sources. The 3rd row is the EEG data calculated from forward model after the source is reconstructed from our proposed model with α = 0.01. The 4th-7th row is the time series plots when α = 0.02, 0.05, 0.1, 0.5 respectively. The topoplots on the right part of Fig.11 is are sampled from 42 ms of the EEG data on the left.

**Figure 10:**
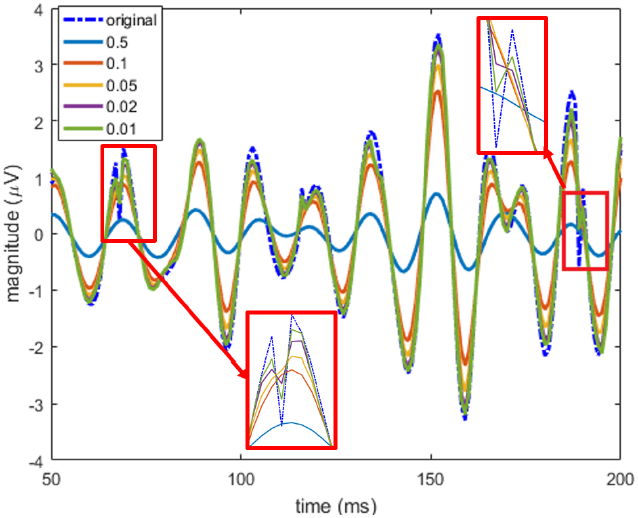
Illustration of the smoothing effect of temporal graph regularization: reconstructed time courses from varied graph regularization parameters.

**Figure 11:**
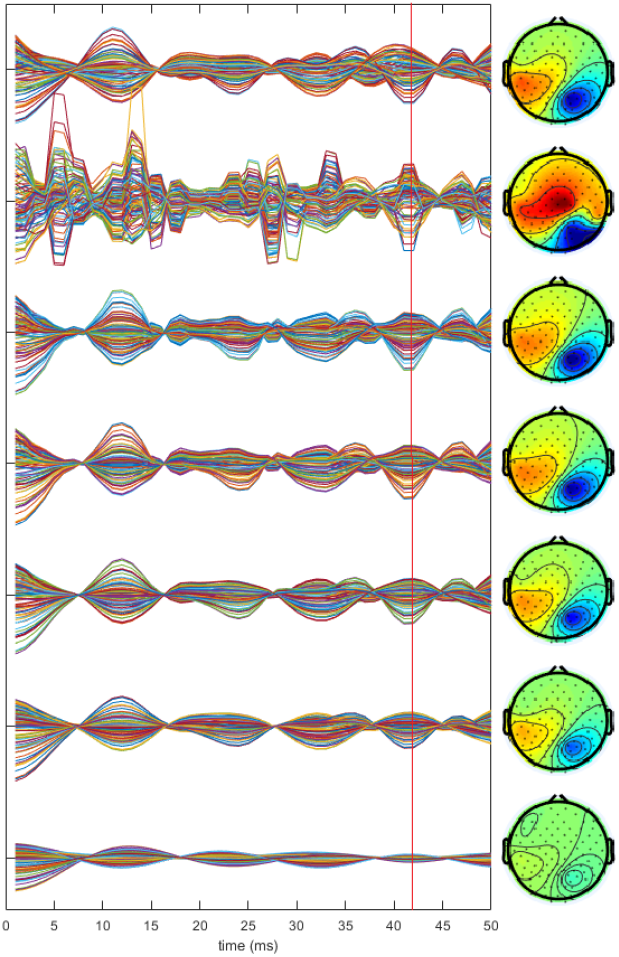
EEG time series plot of the uncorrupted EEG signal, corrupted EEG signal vs the reconstructed EEG signal using the proposed method and the corresponding topoplots at 42 ms: The 1st row is the time series plot of the original uncorrupted EEG data, the 2nd row is the plots for corrupted EEG data, the 3rd row is the EEG data reconstructed by applying our algorithm with α = 0.01, the 4th row is the reconstructed EEG data with α = 0.02, the 5th row is the reconstructed EEG data with α = 0.05, the 6th row is the reconstructed EEG data with α = 0.10, the 7th row is the reconstructed EEG data with α = 0.5. The spurious source in the source space corrupted the task-related EEG data, and by using our graph regularized LRR model, the true EEG data can be recovered.

### 5.4 Experiments 3: Comprehensive Comparison with Benchmark Algorithms

The purpose of previous numerical experiments is to validate each term of the goal function and to understand their properties. The trade-off between low-rankness, data fidelity, sparsity, temporal smooth are fully discussed by varying different parameters. In this part, a comprehensive study is conducted to compare the proposed algorithm with the popular ESI algorithms such as MNE [33], sLORETA [34], MCE and well as the state-of-art algorithm mixed-norm estimate (MxNE) [27].

We generated independent sources in different ROIs for easy validation purpose, the number of independent sources varied from 2 to 4 corresponding to different rank of ground-truth source, and the number of spurious source are generated by randomly activating the sources on the cortex with a random scalar whose mean value to be 0, and the variance is 1. Moreover, the noise on sensor level is also added to the EEG data. Two of the MCE algorithms are selected, which are Homotopy and FISTA [59]. For MxNE algorithm, we choose *l*
_2,1_ to enforce *l*_2_ norm on the temporal time series of each voxel and *l*_1_ norm to impose spatial sparsity.

To measure the performance, we introduced 5 metrics, including 1) CPU time in seconds, 2) rank of the calculated source, 3) Sparsity, measuring the number of nonzero elements in the source space at each time point, 5) Reconstruction Error (RE) defined in Eq.(22), 5) Localization Error (LE), which is calculated using the shortest path algorithm over the irregular meshes from the reconstructed source location to the ground truth location. The LE metric is the most important one, since it measures the discrepancy in location, the other metrics give information of the property of the rendered solution. To calculate LE for each ROI with activated sources, we first locate source with the largest activation magnitude in this ROI, and calculate the shortest path distance from the located source to the ground truth location. We conduct the same procedure for all the activated ROI, and calculate the average value of all the distances at each time point. The final LE is the averaged distance value for all the 500 time points for each experiments.

All the algorithms are implemented in Matlab except MxNE which we call as a Python script (MultiTaskLasso.py) under the linear model of scikit-learn library[60] from Matlab. The formulation for MxNE is

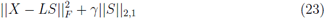

We found that parameter tuning process for MxNE algorithm is very time consuming, even though there is only one parameter for the unweighted version. If the parameter γ in Eq.(23) is tuned for one experiment with good performance, and we use the same setting of parameter and the same setting to generate random source activation patterns, the reconstructed source matrix can be a zero matrix. We tuned the parameter γ according to the best LE performance when the rank is 2 and the number of spurious activated source is 2. By conducting experiments for parameter tuning of MxNE, we set γ in Eq.(23). For our proposed algorithm, we set λ = 0.01 and *β* = 0.01, which were tested to have good performance for the same case when the rank is 2 and the number of spurious activated source is 2, and the graph parameter *α* is also set to be 0.01. 10 experiments were conducted under the same setting and the performance of all the algorithms are summarized in Table 1 to Table 3. The SNR is calculated after the noise signal is generated and it was averaged from 10 experiments under the same experiment setting. As can be seen from the tables, our algorithm is the most accurate to locate the task related activated source. The CPU time of our algorithm is between Homotopy and FISTA algorithm. The running time for Python version of MxNE is much faster than our proposed algorithm, but there are many cases that MxNE algorithm failed, thus making the overall accuracy drop significantly. Although our algorithm have more parameter, it is a more sophisticated model that allows better controllability to customize the weight of different terms in the goal function. Although MxNE has large RE and LE, it is still worth noting that the MxNE algorithm we used is a simple version with *ℓ*_2,1_ norm discussed in Ref.[27], and the algorithm to solve the goal function 23 is coordinate descent, other algorithms can be tested to solve the same problem. MCE model solve using *ℓ*_1_ algorithms Homotopy and FISTA can render good LE accuracy, and Homotopy outperforms FISTA in all the experiment with better speed, which confirms the comparison discussed in [40][56].

**Table 1:** Source Reconstruction Performance Comparison (Rank=2)

| Method | Rank =2; SNR= -0.356 dB |  |  |  |  | Rank =2; SNR= -1.12 dB |  |  |  |  | Rank =2; SNR= -1.67 dB |  |  |  |  |
| --- | --- | --- | --- | --- | --- | --- | --- | --- | --- | --- | --- | --- | --- | --- | --- |
|  | Time | Rank | Sparsity | RE | LE | Time | Rank | Sparsity | RE | LE | Time | Rank | Sparsity | RE | LE |
| Homotopy | 0.41 | 449 | 65 | 0.80 | 2.02 | 0.41 | 449.8 | 84.4 | 0.95 | 3.31 | 0.50 | 459.3 | 91.15 | 0.97 | 4.57 |
| FISTA | 2.76 | 500 | 1812.4 | 0.62 | 3.93 | 2.76 | 500 | 1920.4 | 0.89 | 9.39 | 2.73 | 500 | 1888.0 | 0.80 | 11.25 |
| sLORETA | 0.055 | 500 | 2004 | 1.18 | 33.5 | 0.05 | 500 | 2004 | 1.26 | 34.13 | 0.056 | 500 | 2004.0 | 1.32 | 38.31 |
| WMN | 8.4e-5 | 107 | 2004 | 0.99 | 22.6 | 8.4e-5 | 107 | 2004 | 0.99 | 31.47 | 8.4e-5 | 107 | 2004.0 | 0.99 | 28.46 |
| MxNE | 0.026 | 7.9 | 7.9 | 0.95 | 42.1 | 0.023 | 10 | 10 | 0.97 | 45.79 | 0.028 | 11.8 | 11.8 | 1.04 | 68.80 |
| Proposed | 0.75 | 3.3 | 3.3 | 0.24 | **0.145** | 0.75 | 4.8 | 4.8 | 0.414 | **1.94** | 0.83 | 6.4 | 6.5 | 0.42 | **2.27** |

**Table 2:** Source Reconstruction Performance Comparison (Rank=3)

| Method | Rank =3; SNR= 0.938 dB |  |  |  |  | Rank =3; SNR= -0.174 dB |  |  |  |  | Rank =3; SNR= -0.784 dB |  |  |  |  |
| --- | --- | --- | --- | --- | --- | --- | --- | --- | --- | --- | --- | --- | --- | --- | --- |
|  | Time | Rank | Sparsity | RE | LE | Time | Rank | Sparsity | RE | LE | Time | Rank | Sparsity | RE | LE |
| Homotopy | 0.36 | 445.4 | 77.6 | 0.64 | 5.18 | 0.48 | 464 | 87.2 | 0.68 | 6.1 | 0.46 | 474.1 | 92.5 | 0.84 | 5.88 |
| FISTA | 2.64 | 500 | 1812.5 | 0.69 | 9.33 | 2.56 | 500 | 1969.0 | 1.23 | 21.5 | 2.67 | 500 | 1939.2 | 0.94 | 17.44 |
| sLORETA | 0.057 | 500 | 2004 | 1.90 | 41.93 | 0.06 | 500 | 2004 | 1.81 | 43.9 | 0.05 | 500 | 2004.0 | 1.25 | 46.60 |
| WMN | 8.4e-5 | 107 | 2004 | 6.79 | 29.55 | 7.6e-5 | 107 | 2004 | 6.82 | 35.63 | 7.3e-5 | 107 | 2004.0 | 0.99 | 34.41 |
| MxNE | 0.026 | 9.2 | 9.2 | 0.91 | 64.95 | 0.028 | 10 | 10 | 0.98 | 67.95 | 0.028 | 12.2 | 12.2 | 0.96 | 57.49 |
| Proposed | 0.72 | 5.2 | 5.2 | 0.43 | **2.98** | 0.80 | 6.1 | 6.1 | 0.488 | **3.82** | 0.75 | 7.3 | 7.3 | 0.45 | **2.86** |

**Table 3:** Source Reconstruction Performance Comparison (Rank=4)

| Method | Rank = 4; SNR= 1.04 dB |  |  |  |  | Rank =4; SNR= 0.603 dB |  |  |  |  | Rank =4; SNR= -0.179 dB |  |  |  |  |
| --- | --- | --- | --- | --- | --- | --- | --- | --- | --- | --- | --- | --- | --- | --- | --- |
|  | Time | Rank | Sparsity | RE | LE | Time | Rank | Sparsity | RE | LE | Time | Rank | Sparsity | RE | LE |
| Homotopy | 0.45 | 448 | 78.7 | 0.57 | 3.77 | 0.41 | 467.3 | 87.9 | 0.62 | 5.87 | 0.46 | 475.1 | 93.15 | 0.62 | 5.35 |
| FISTA | 2.97 | 500 | 1978.7 | 1.03 | 16.54 | 2.46 | 500 | 1964.5 | 1.01 | 16.64 | 2.45 | 500 | 1961.9 | 1.35 | 19.4 |
| sLORETA | 0.058 | 500 | 2004 | 1.99 | 45.3 | 0.058 | 500 | 2004 | 1.72 | 49.26 | 0.050 | 500 | 2004.0 | 1.71 | 49.38 |
| WMN | 7.8e-5 | 107 | 2004 | 7.43 | 31.0 | 7.8e-5 | 107 | 2004 | 6.63 | 33.49 | 7.8e-5 | 107 | 2004.0 | 6.66 | 38.00 |
| MxNE | 0.024 | 11.9 | 11.9 | 0.997 | 68.36 | 0.023 | 11.7 | 11.7 | 0.947 | 56.42 | 0.025 | 13.9 | 13.9 | 0.99 | 61.46 |
| Proposed | 0.79 | 5.9 | 6.8 | 0.38 | **2.24** | 6.9 | 7.8 | 7.8 | 0.43 | **3.90** | 0.76 | 10.1 | 14.3 | 0.44 | **4.5** |

## 6. Conclusion

In this paper, we propose to consider the noise not only on the sensor level, but also in the source space. we come up with an EEG source imaging model based on temporal graph structures and low-rank representation. The model is solved with our proposed algorithm based on ADMM. Numerical experiments are conducted to verify the effectiveness of the proposed work on discovering task related low-rank sources. We delineate the discussion on the properties and impacts for each term in the cost function to help better understand our proposed graph regularized low rank representation model. Compared the traditional model, our proposed one can find the intrinsic task related activation patterns and suppress the spurious source patterns.

## 7. Acknowledgment

This work been partially supported by the NSF funding under grant number CMMI-1537504 and DMS-1522786. The authors are also grateful to Dr. Stefan Haufe and Dr. Yu Huang for providing the head model.

